# Avoiding misinterpretation of regression lines in allometry: is sexual dimorphism in digit ratio spurious?

**DOI:** 10.1101/298786

**Authors:** Wolfgang Forstmeier

## Abstract

The statistical analysis of allometry (size-dependence of traits) is fraught with difficulty that is often underestimated. In light of some recent controversies about statistical methods and the resulting biological conclusions, I here discuss the interpretation of regression lines and show how to avoid spurious effects. General linear models based on ordinary least square (OLS) regression are often used to quantify sexual dimorphism in a trait of interest that is modelled as a function of sex while controlling for size as a covariate. However, an analysis of artificially generated data where males and females differ in size only, but are otherwise built according to the same principles, shows that the OLS method induces a spurious dimorphism where there is none. Hence, OLS-based general linear models should not be regarded as a fail-proof tool that automatically provides the correct answer to whatever question one has in mind. Here I show how to avoid misinterpretation and how to best proceed with answering the recent debate about sexual dimorphism in digit ratio, a trait that is thought to reflect sex-hormone levels during development. The limited data, currently available to me, suggests that the widely accepted sexual dimorphism in digit ratio might well be only a by-product of an allometric shift in shape, urgently calling for a re-examination in larger data sets on humans and other vertebrates.

## A controversy about sexual dimorphism as an artefact

The relative length of human index and ring fingers (digit ratio, 2D:4D) has received a lot of research attention, because it is known to be sexually dimorphic (smaller ratios in men than in women [1, 2]) and has been suggested to reflect sex-hormone levels experienced during development [2–5]. Now Lolli and coworkers [6] have made a claim that, if true, would have major consequences for this research field. According to their analyses of data on human finger lengths, the apparently lower average digit ratio of men compared to women arises simply as an artefact of allometry: as hands get bigger, there is a general shift in shape such that digit ratio gets smaller. In consequence, there is no particular sexual dimorphism (apart from men being generally larger) that would require an additional explanation invoking specific sex-hormone effects on finger development.

This potentially controversial conclusion might however be questioned for methodological reasons. The study by Lolli and coworkers [6] investigates allometry and sexual dimorphism by regressing the length of the second finger 2D (dependent variable) over the length of the fourth finger 4D (predictor variable), yet the implemented method of ordinary least square (OLS) regression fails to account for ‘biological noise’ (aka ‘natural variation’ [7] or ‘biological deviance’ [8]) in the predictor variable. An earlier paper [9] already analysed the very same data set using such OLS regression and was subsequently criticized for its inappropriate statistical approach [10]. Indeed, it is easy to show (see further below) that OLS regression systematically induces positive sexual dimorphism (relatively larger trait values in the larger sex) and is therefore not suitable for quantifying sexual dimorphism [10, 11]. Yet, Lolli and coworkers [6] did not address these statistical concerns but rather followed the earlier study [9] (and others, e.g. [12]) in using OLS-based methods. Apparently this was also motivated by a recent methodological review by Kilmer and Rodríguez [13], who incautiously recommend OLS regression for studies of allometry, despite of the inherent problems [7, 10, 14, 15] that have caused a long-lasting debate about regression methods [7, 8, 10]. This may lead to the unfortunate situation where readers are generally confused about which method of regression to use. To bring more clarity into this debate, I here would like to highlight two points:

(I) The interpretation of OLS regression lines in the study of allometry and sexual dimorphism is more difficult than is typically recognized [6, 13]. OLS regression lines are suitable for predicting values of y from a given value of x, but they are not designed to automatically reflect the principles that underlie allometric relationships [7]. If we aim at finding out whether males and females differ in size only but are otherwise built according to the same general principles, then OLS regression lines will typically give the wrong answer. For correct interpretation of sexual dimorphism ‘after accounting for differences in size’, regression slopes need to be adjusted for biological noise in the predictor variable. In contrast, accounting only for technical measurement error in the predictor variable (as suggested by Kilmer and Rodríguez [13]) is not sufficient for removing the spurious dimorphism.

(II) I suggest a different way of how to analyse whether digit ratio is sexually dimorphic after accounting for body size. Rather than plotting one measure of size over another (2D over 4D, which calls for major axis regression [7, 10, 15, 16]), I suggest to plot digit ratio directly over mean finger length, and to adjust OLS regression slopes for the biological noise in the predictor. Intriguingly, such analysis of the empirical data made available by Lolli and coworkers [6] indeed suggests that sexual dimorphism in digit ratio may simply emerge from an allometric change in shape with size.

Ratios are widely used in many disciplines (e.g. body mass index, waist to hip ratio, body condition index) because many ratios have advantageous properties, which however rarely get appreciated explicitly [15]. More frequently, the inconsiderate use of ratios has been criticized [16–19], typically because ratios often fail to achieve independence of body size. By their very nature, ratios (here 2D:4D) are strongly positively correlated with their numerator (here 2D) and strongly negatively correlated with their denominator (here 4D), yet these are mathematical necessities that do not reveal whether a ratio is size dependent. Ratios can in principle be independent of variation in body size (see below), but whether real data on human digit ratios show such independence of size (here average finger length) is a matter of empirical testing.

## Does digit ratio shift continuously with size?

Figure 1a shows digit ratios calculated from the data of Lolli and coworkers [6] plotted over the mean length of the two fingers. Intriguingly, within each sex, digit ratio declines as fingers get longer, and the two OLS regression lines for males and females (the appropriateness of which I will address below) practically coincide. This suggests that, as fingers get longer on average, the ratio 2D:4D is shifting continuously, with 4D getting longer disproportionally and 2D lagging behind. Judging from the indicated OLS regression lines, there is apparently no need to invoke any sexual dimorphism in digit ratio beyond the allometric shift. This is because at the population-wide average finger length of 70.4 mm, men and women are predicted to have practically equal digit ratios (0.9890 and 0.9895 respectively), while without consideration of the allometric shift, the sexes are clearly different in their average digit ratio (men: 0.9828, women: 0.9925, t_830_ = 3.68, p = 0.0003). Whether the data shown in Figure 1a is truly representative of human allometry will have to be shown in the future by additional analyses of larger data sets, but it seems possible that the phenomenon is more general.

**Figure 1.**
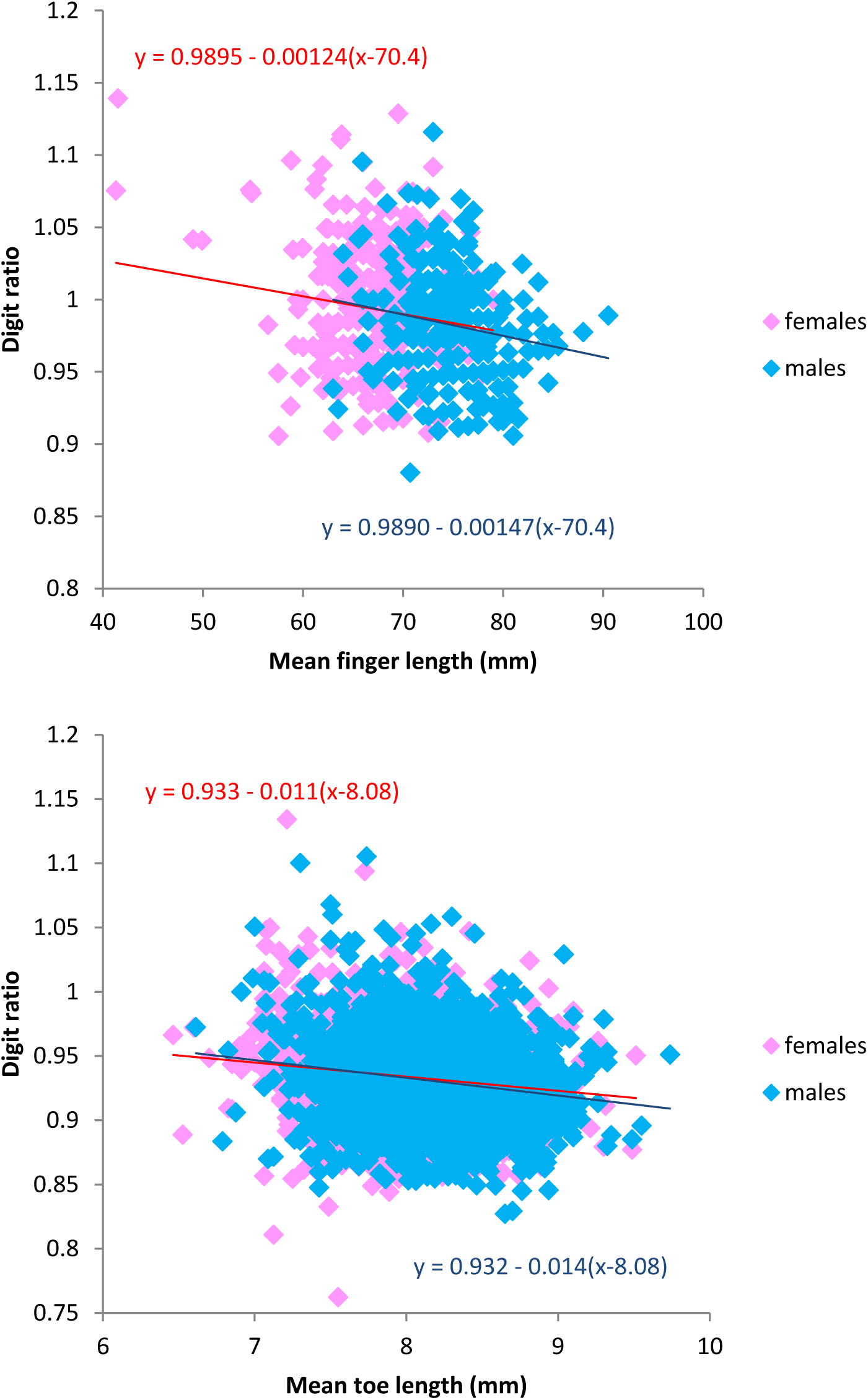
A: Digit ratios (2D:4D) of men and women in relation to mean finger length ((2D + 4D) / 2) calculated from the data provided by [6], both left and right hands analysed jointly. OLS regression lines are indicated for each sex. B: Digit ratios (2D:4D) from the right foot of male and female zebra finches in relation to mean toe length ((2D + 4D) / 2). Sex-specific OLS regression lines are shown. Intercepts are reported at the population mean by subtracting the population mean from the covariate.

The human digit ratio literature has also inspired similar research on numerous other tetrapod species, many of which have also been found to be sexually dimorphic for digit ratio (e.g. [20–27]). Hence, I examined own data on digit ratio in zebra finches (2D:4D, based on toes of the right foot; see [28]). Figure 1b illustrates an interesting similarity with the human case, since within both males and females digit ratio continuously declines as digits get longer and the two OLS regression lines practically coincide. Since the toes of males are only slightly longer than those of females (8.14 mm versus 8.02 mm) and the regression line is relatively shallow, male digit ratios are shifted to only slightly smaller values (mean 0.9310) compared to those of females (mean 0.9336). This sexual dimorphism in digit ratio only barely reaches statistical significance p = 0.04 despite a sample size of n = 3389, and of course disappears when accounting for the allometric shift. This illustrates further the need to reassess the presence of sexual dimorphism in digit ratio beyond what can be explained by allometry [26], not only in humans but also in other species [20–25, 27].

Previously [10], I argued that OLS regression can lead to erroneous conclusions about sexual dimorphism because it fails to acknowledge that there is biological noise in the predictor variable (e.g. when regressing 2D over 4D). If our interest lies in exploring sexual dimorphism, we should consider major axis regression (MA) or reduced major axis regression (RMA) when regressing one measure of size over another, because these methods assume equal amounts of statistical noise in the two variables, which appears most sensible for two digits with comparable properties [10]. Now in the above case (Fig. 1), I used OLS regression merely out of convenience, as is often done, but is this really appropriate, given that the x-axis of Figure 1 contains biological noise? Below I will show with simulated data that, in some sense, the regression lines in Figure 1 are indeed somewhat too shallow, being biased downwards by the biological noise in the x-axis. The slopes should in fact be adjusted (see [29]) if, and only if, the aim of line-fitting is to understand the general principles from which males and females are built.

## Creating simulated data that include biological noise

Closely adhering to the simulations that I carried out previously [10], I created artificial data on finger lengths following specific predefined rules. In the first scenario, I assume isometry (like in [10]) simply to illustrate how one gets to the situation where ratios are size independent. In the second scenario, I incorporate an allometric shift with size (importantly, the same for the two sexes), and then examine which regression method allows retrieving the correct value for the slope of the underlying line around which the data have been generated. The data generation always consists of three steps: (1) generating between-individual variation in latent size of fingers (representing the sum of genetic and environmental effects that make the fingers of some individuals generally shorter or longer, thereby inducing a strong positive correlation between 2D and 4D across individuals, r = 0.9 in [6]), (2) applying underlying rules of isometry or allometric shift, and (3) adding individual biological noise to each finger separately (representing the sum of unique genetic and environmental effects on each finger, which induces the scatter around the regression lines in plots of 2D over 4D, and which is the main source of individual differences in digit ratio).

For the first scenario of isometry, latent finger lengths of 5,000 women were drawn from a normal distribution with a mean of 69 mm and a standard deviation (SD) of 5 mm. For 5,000 men I used a mean of 77 mm and also a SD of 5 mm. Next, for women I assumed perfect isometry with a slope of 1 (2D = 4D = latent size), corresponding to a digit ratio of 1. For men, I assumed an isometric slope of 0.985 (2D = 0.985*4D = 0.985* latent size), corresponding to a digit ratio of 0.985. Finally, for each finger I drew biological noise from a normal distribution with a mean of zero and a SD of 2 mm. Yet to reflect the fact that the amount of noise is typically proportional to size, I multiplied each value of noise with the relative latent size (i.e. with latent size / 73 mm). This means that the noise was on average 10% larger for an individual whose latent size was 10% above the population mean. After adding the created noise to each finger separately, I calculated for each individual its realized mean finger length ((2D + 4D) / 2) and its digit ratio (2D / 4D).

In the second scenario, latent sizes were created as above. Yet this time digit ratio was not assumed to be size-independent, but (inspired by Figure 1a) rather to change systematically with latent size, equally in the two sexes (expected digit ratio = 0.984 – 0.002 * (latent size – 73)). Using this equation, I calculated expected digit ratios for each individual from its latent size. The expected length of 2D was then determined as 2D = (2 * latent size * expected digit ratio) / (1 + expected digit ratio), and the expected length of 4D as 4D = 2D / expected digit ratio. After this, I again added biological noise to each finger as described above and calculated realized mean finger length and digit ratio.

## OLS regression does not automatically retrieve the underlying principles

Figure 2a illustrates the first scenario of isometry, where digit ratios are independent of size, and where sexual dimorphism in digit ratio can be judged from the sex-specific intercepts at the population mean size of 73 mm.

**Figure 2.**
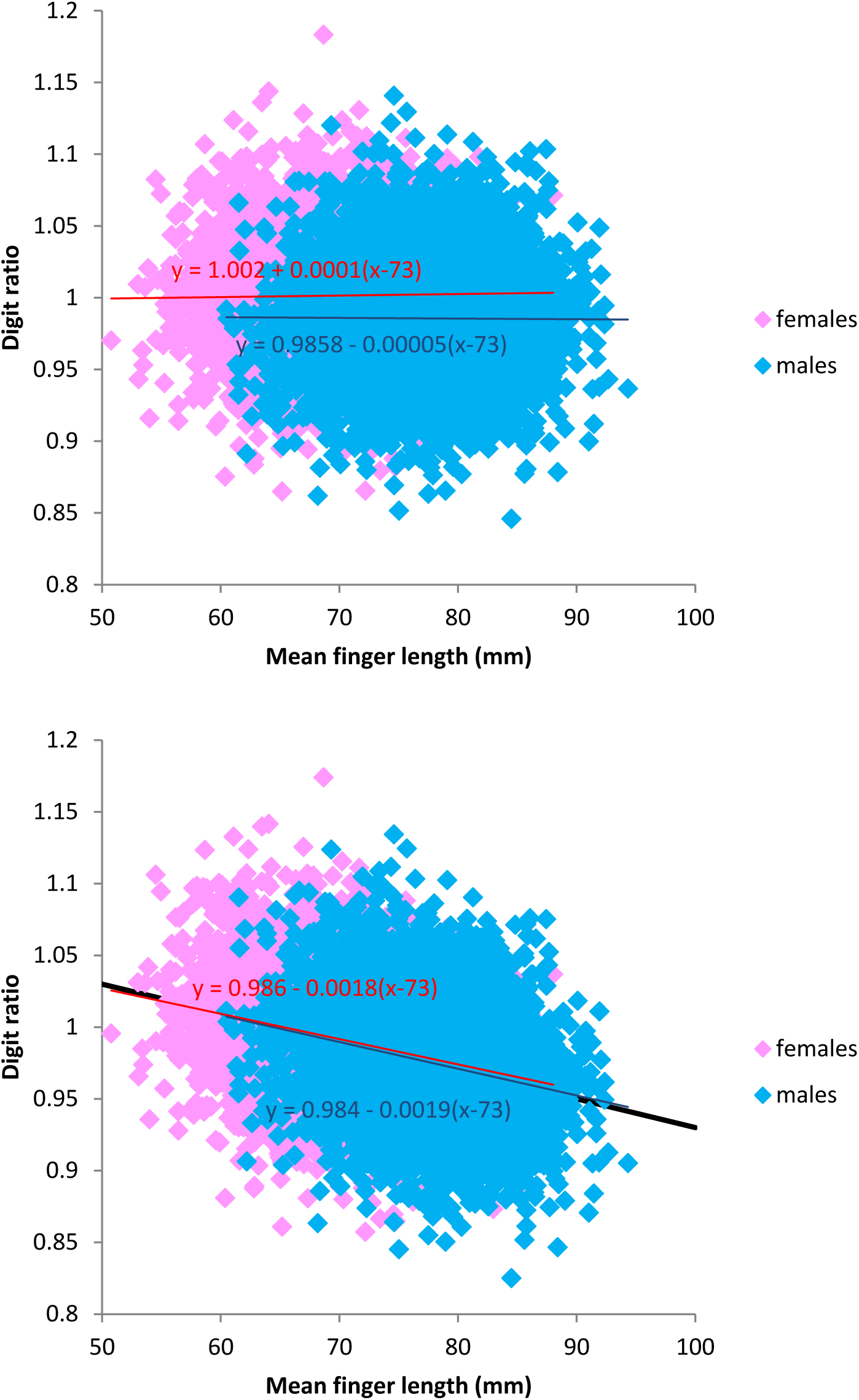
Digit ratio in relation to mean finger length calculated from simulated data on 5,000 individuals of each sex, together with sex-specific OLS-regression lines. A: Case of isometry (size independence) combined with sexual dimorphism in digit ratio. B: Case of an allometric shift in digit ratio with size (mean finger length) that is shared between the sexes. Note that the thick black line in the background (y = 0.984 -0.002(x-73)) shows the underlying change in digit ratio with latent size rather than with realized mean finger length.

Figure 2b illustrates the second scenario of allometric shift where digit ratios are size dependent, together with sex-specific OLS regression lines. The obtained OLS regression slopes for 5,000 males (-0.0019, thin blue line in Fig. 2b) and 5,000 females (-0.0018, red line) were marginally shallower than the slope of the line around which the data were generated (-0.0020, thick black line). Given this marginal difference in slopes which could have arisen by chance from sampling noise, I repeated the simulation based on 100,000 individuals of each sex and this was replicated 20 times to calculate a between-replicate standard error of slope estimates, each obtained from 100,000 individuals. Indeed, the average slope for males (-0.00182 ± 0.00001) and females (-0.00187 ± 0.00001) are clearly shallower than the slope of the underlying digit ratio change with latent size (-0.00200).

The reason behind this attenuation in slopes lies in the fact that the x-axis of Figure 2b shows the realized size (after adding biological noise to each finger), while the slope of the underlying relationship (-0.002, black line) refers to the change with latent size (before adding the noise). Which of these slopes is considered as ‘correct’ is an issue of the purpose of line fitting. The shallower OLS regression lines represent the optimal solution for predicting for each sex an individual’s digit ratio from its realized mean finger length. However, these sex-specific lines are too shallow if one wants to retrieve the rules according to which the data had been generated (namely a universal rule that is shared between the sexes rather than two sex-specific rules).

Importantly, the shallower OLS lines in Figure 2b are not “flawed” since the x-axis of Figure 2b shows the realized mean finger length rather than latent size, but they can be misinterpreted. When researchers say “after accounting for variation in size” they often forget that they have accounted for ‘realized size’ but not for ‘latent size’ (which remains unknown in real data sets) and hence they cannot retrieve the rules from which the sexes were built.

The degree of attenuation of slopes caused by the biological noise in the x-axis is relatively straightforward to calculate for such simulated data (see [29]). For each sex, the between-individual variance in latent size is 25 mm^2^ (the square of the SD (= 5 mm) of the normal distribution from which latent size was sampled). To each finger we added noise with a variance of 4 mm^2^ (SD = 2 mm). Since the realized average finger length is calculated from two fingers ((D4 + D2) / 2), the noise in the averages is half of that in each finger (4 mm^2^ / 2 = 2 mm^2^). Given the additivity of variances, the between-individual variance in realized mean finger length is 25 mm^2^ + 2 mm^2^ = 27 mm^2^. Yet since the amount of noise was scaled to latent size, there is overall 25 mm^2^ + 2.2 mm^2^ = 27.2 mm^2^ for males and 25 mm^2^ + 1.8 mm^2^ = 26.8 mm^2^ for females. The ratio of variance in latent size to variance in realized size (25/27 = 0.926) quantifies the amount of attenuation of slopes (0.918 for males and 0.933 for females). Dividing the obtained OLS regression slopes from Figure 2b by the attenuation factor, retrieves the slope of the underlying relationship with latent size (-0.002) which was the same by design for both sexes (males: -0.00182 / 0.918 = -0.002; females: -0.00187 / 0.933 = -0.002).

## Careful interpretation of regression lines is needed

The above simulation illustrates that OLS-based general linear models should not be regarded as a fail-proof tool that automatically provides the correct answer to whatever question one has in mind. If we set up a scenario where the sexes only differ in their size but otherwise are constructed according to identical rules, we would ideally like to be able to retrieve that information from our data analysis. Specifically, after correcting for variation in size, we would like to see identical intercepts for the two sexes. This was not the case here, because OLS slopes were systematically biased downwards as a function of the amount of biological noise in the x-axis. In the present case of Figures 1 and 2, the amount of downward bias in slopes is almost negligible (here about 7-8 %), such that the resulting sexual dimorphism (women having slightly higher digit ratios than men at the population average finger length of 73 mm, see Fig. 2b) is even difficult to detect. Here, OLS regression fails to account for the biological noise in the x-axis, but the resulting difference in intercepts is really minimal (though statistically significant, see final paragraph). The resulting bias is minimal because, in this example, the y-axis contains much more biological noise than the x-axis (here making OLS regression much more sensible than RMA regression).

This situation clearly changes when examining apparent sexual dimorphism from OLS regression lines in plots of one measure of size over another. To illustrate that point, I plotted the data from Figure 2b in such a way: once arbitrarily plotting 2D over 4D (Fig. 3a) and once arbitrarily plotting 4D over 2D (Fig. 3b). The OLS regression slopes show the expected bias, namely an apparent (deceptive) sexual dimorphism with the larger sex (males) showing the ‘relatively’ larger trait values (irrespective of what the trait is). The lines suggest that males have relatively longer 2D than females (by 0.85 mm = 72.98 mm – 72.13 mm) after accounting for variation in 4D (Fig. 3a), and when plotting the very same data the other way around, the lines suggest that males have relatively longer 4D than females (by 1.50 mm = 74.50 mm – 73.00 mm) after accounting for variation in 2D (Fig. 3b). The fact that these two interpretations appear paradoxical (it is confusing when both fingers are relatively longer in males after accounting for size), illustrates that it is erroneous to interpret OLS regression lines in such a way (see [7, 10, 11]).

Hence, OLS regressions should be interpreted with greatest care, and are not an easy tool in the study of allometry. Prominent claims to the contrary [13] are based on the misunderstanding that biological noise could be ignored when measurement error is accounted for. Measurement error is a minor technical problem that can be effectively reduced by averaging multiple repeated measures of an individual’s phenotype. Such measurement error is not the primary reason why individual data points in Figure 3 deviate from the regression lines [7, 8, 29]. The main reason for deviation from the line lies in biological noise. Biological noise, in the present example, is the sum of genetic and environmental effects that let an individual’s phenotype deviate from its ‘expected value according to its latent size’ (e.g. alleles that affect one finger but not the other) [8]. This biological noise is likely of about equal magnitude for 2D and 4D, so it makes little sense to declare one of them to be the dependent variable and the other one the predictor. In such symmetrical problems MA or RMA regression is most helpful [7, 8, 29]. Surely OLS regression can be used in plots of one measure of size over another (e.g. brain mass over body height; [30]) but the obtained regression lines should never be interpreted as proof of sexual dimorphism in y after accounting for variation in x (e.g. the erroneous claim that men have relatively larger brains than women after correcting for body size, [10, 30]). Given that the explicit modelling of biological noise in the x-axis is still far from commonplace, one should always be maximally cautious with formulations like ‘after correcting for differences in body size…’.

**Figure 3.**
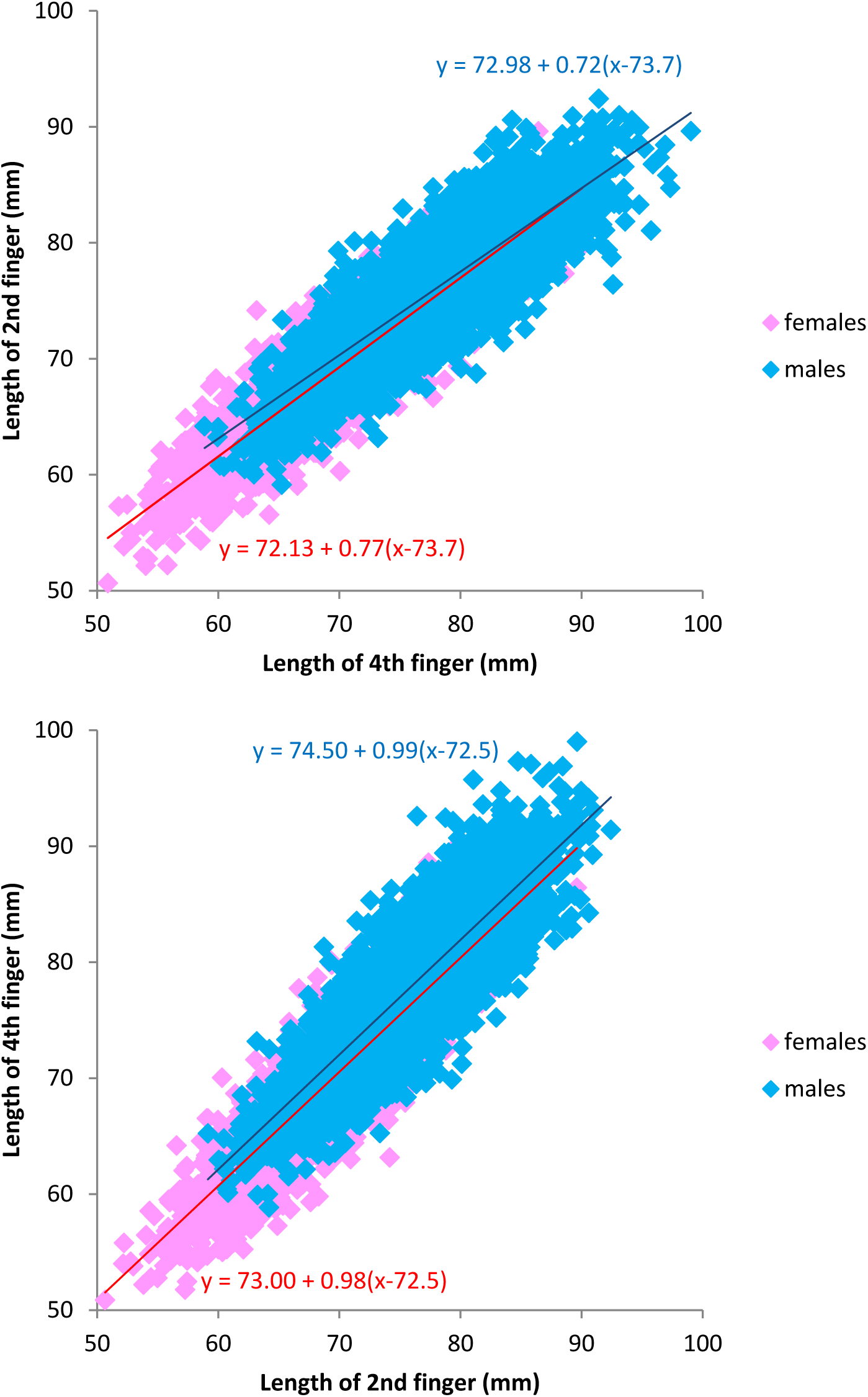
The data that was generated for Figure 2b, plotted in a way where OLS-regression lines are particularly confusing because in each case the blue line for males lies above the red line for females. A: Plot of 2D over 4D with sex-specific OLS-regression lines suggesting that, at the population mean of 4D of 73.7 mm, males have slightly longer 2D (72.98 mm) than females (72.13 mm) after accounting for the variation in 4D. B: Plot of 4D over 2D with sex-specific OLS-regression lines suggesting that, at the population mean of 2D of 72.5 mm, males have longer 4D (74.50 mm) than females (73.00 mm) after accounting for the variation in 2D.

## Next steps for digit ratio research

For the interpretation of digit ratio as a marker of sex-hormone effects, it appears essential to know whether digit ratio is sexually dimorphic or not once accounting for any size-related shifts like in Figure 1 (see also [26, 27]). This question should be relatively easy to answer, given that numerous studies have already collected the relevant morphological measurements. Assuming that most researchers are unfamiliar with Bayesian modelling of variation in latent size, what is the most pragmatic way of distinguishing between the two scenarios illustrated in Figure 2a versus Figure 2b?

I recommend analysing digit ratio as the dependent variable in a general linear model (effectively an ANCOVA) with sex as a fixed effect and mean finger length as a covariate (main effects without interaction), and interpreting the model output carefully with the attenuation in OLS regression slopes in mind. Such an analysis of the data shown in Figure 2a, yields a highly significant main effect of sex (estimate -0.016 ± 0.001, t = -16.3, p < 10^-16^, note that n = 10,000) and a non-significant slope estimate for the covariate ‘mean finger length’ (estimate 0.00003 ± 0.00007, t = 0.4, p = 0.7). Essentially, we want to verify that there is not an overall negative regression slope (like in Figure 2b) that runs through the average coordinates of the male data and the average coordinates of the female data. The slope of such a hypothetical line can be calculated as (average male digit ratio – average female digit ratio) / (average male finger length – average female finger length) = -0.00200. The lower end of the 95% CI of the model’s slope estimate (0.00003 – 1.96*0.00007 = -0.00012) lies still clearly above that value, even when considering that there is some attenuation of the OLS regression slope.

For comparison, the analysis of the data from Figure 2b, yields a small and only barely significant main effect of sex (estimate -0.002 ± 0.001, t = -2.3, p = 0.02) and a highly significant negative slope for the covariate ‘mean finger length’ (estimate -0.0018 ± 0.00007, t = -24.4, p < 10^-16^). Thanks to the very large sample size, the artefacts of spurious dimorphism and attenuation of slopes become visible. Although it seems like there is still a small but significant effect of sex, and also the lower end of the 95% CI of the slope estimate (-0.00196) is only barely above the hypothetical line (-0.00200), these minor deviations should be attributed to the attenuation of the OLS regression slope. How much attenuation should be allowed for? To find out empirically, I modified the parameters of the data simulation shown in Figure 2b until it most closely resembled Figure 1a (when using SD of latent size = 4.2 and SD of noise = 1.85). With these parameters the attenuation of slopes is about 10% (4.2^2^/ (4.2^2^ + 0.5*1.85^2^) = 0.91), so slope estimates should be multiplied by the inverse of that (1 / 0.91 = 1.097). Generally, the lower the Pearson correlation between 2D and 4D across both sexes (e.g. r = 0.9 in Figure 3), the higher the noise component and the stronger the attenuation of slopes in our analysis of Figure 2b.

Several studies (e.g. [20, 21, 26]) have assessed the size-dependence of digit ratio by plotting it over other measures of size than mean finger length (e.g. body mass), which bears a risk of overlooking a size dependence because the covariate might contain additional biological noise that is not relevant for digit ratio. Hence regressing digit ratio over mean finger length appears most effective in identifying a direct dependency.

Sample size is important to consider, if our aim is to reject the idea that there is an allometric shift in digit ratio like shown in Figure 1a. In this analysis, I have treated measurements from both hands as equivalent in order to maximise statistical power. Generally, I would recommend running separate analyses for each hand, only if the patterns differ significantly between hands. In Figure 1a, the Pearson correlation coefficient between digit ratio and mean finger length (across both sexes) is about r = -0.20. A total sample size of about 260 individuals is required to achieve a power of 95% for detecting such a correlation (with α = 0.05; calculated using G*Power 3.1.7, [31]). If one aims at detecting correlations even within each of the two sexes (here r = -0.15 for females and r = -0.17 for males), a sample size of about 470 individuals per sex would be required for a statistical power of 95%. Studies with a lower statistical power (e.g. [32]) should still report the overall correlation between digit ratio and mean finger length, for a later meta-analytic summary.

If the patterns shown in Figure 1 should turn out to be representative of humans and also other animals, it is not easy to see how digit ratio could be a more informative indicator of sex-hormone effects than for instance mean digit length. Ignoring the allometric shift, the sex difference in human digit ratio is only about 0.26 within-sex standard deviations (Cohen’s D calculated from Figure 1a), while the sex difference in mean finger length is about six times larger (Cohen’s D = 1.56, Figure 1a), hence the latter somehow appears to have greater potential in reflecting levels of sex hormones experienced during development.

Proponents of the idea that digit ratio is informative about sex-hormone effects face a two-fold challenge. First, they should demonstrate that sexual dimorphism in digit ratio exists independently of the dimorphism in size (Figure 2a versus Figure 2b), and that the trait is more informative than other sexually dimorphic characters. Second, when testing for the allometric shift, they should demonstrate the objectivity of their analyses in spite of a possible conflict of interest or at least some wishful thinking. To show that one has not e.g. selectively removed outliers that appear influential after inspecting the data (see top left corner of Fig. 1a), I suggest to first deposit complete data sets that have been used in a previous publication (that allow reconstructing the results shown in that publication), and then to analyse the complete data set, following the approach outlined above. This effectively eliminates most ‘researcher degrees-of-freedom’ [33] and thereby demonstrates a maximum of scientific objectivity [34].

## Acknowledgements

I thank Stefan van Dongen, Pim Edelaar and Jarrod Hadfield for valuable comments.

